# *APOE* genotypes differentially remodel the astrocytic lipid droplet-associated proteome to shape lipid droplet dynamics

**DOI:** 10.1101/2025.08.19.669163

**Authors:** Carla Cuní-López, Jessica T. Root, Ying Hao, Isabelle Kowal, Niek Blomberg, Rodolfo Ghirlando, Linda G. Yang, Sascha J. Koppes-den Hertog, Mark R. Cookson, Rik van der Kant, Martin Giera, Yue Andy Qi, Priyanka S. Narayan

**Affiliations:** Genetics and Biochemistry Branch, National Institute of Diabetes and Digestive and Kidney Diseases, National Institutes of Health, Bethesda, MD USA; Center for Alzheimer’s and Related Dementias (CARD), National Institutes of Health, Bethesda, MD, USA; Leiden University Medical Center, Center for Proteomics and Metabolomics, Leiden, the Netherlands; Laboratory of Molecular Biology, National Institute of Diabetes, Digestive and Kidney Diseases, National Institutes of Health, Bethesda, MD USA; Center for Neurogenomics and Cognitive Research, Vrije Universiteit Amsterdam, Amsterdam Neuroscience, Amsterdam, the Netherlands; Alzheimer Center Amsterdam, Department of Neurology, Amsterdam University Medical Center, Amsterdam Neuroscience, Amsterdam, the Netherlands; Laboratory of Neurogenetics, National Institute on Aging, National Institutes of Health, Bethesda, MD, USA

## Abstract

The lipids and proteins that comprise lipid droplets regulate several cellular functions including lipid storage, stress responses, and inflammation. Glial lipid droplets have been implicated in the pathogenesis and progression of Alzheimer’s disease (AD), yet the mechanisms linking genetic risk to lipid droplet biology remain unclear. Here we examined how *APOE*, the strongest genetic modulator of late-onset AD, impacts lipid droplet composition and dynamics. We defined the lipid droplet-associated proteome and lipidome in human induced pluripotent stem cell-derived astrocytes harboring the three common *APOE* genotypes: *APOE2* (protective), *APOE3* (neutral), and *APOE4* (risk). Each *APOE* variant displays distinct lipid droplet-associated proteins and lipids. These molecular changes yield differences in lipophagy; lipid droplets in *APOE2* astrocytes undergo autophagic turnover, whereas those in *APOE4* astrocytes are resistant to degradation. These findings suggest that impaired lipid droplet clearance, rather than accumulation, distinguishes *APOE4*-associated AD risk, and may present a new metabolic node for modulating risk.

## Introduction

Lipid droplets are dynamic and understudied organelles composed of a neutral lipid core surrounded by a phospholipid monolayer functionalized with many proteins^1^. Traditionally associated with lipid storage, lipid droplets have roles in several other cellular processes including transcriptional regulation, inflammation, and stress responses^2^. These functions are mediated not only by the lipid content of these organelles but also by the proteins decorating their membranes. Importantly, both cell type and cellular environment modulate the lipid droplet proteome and lipidome^3–9^.

Lipid droplets have been studied extensively in the context of hepatic and adipose tissue, however, these organelles also play key roles in brain cell types. Altered lipid droplet biology has been linked to both brain aging and in neurodegenerative disease^10–20^. Notably, in the first case report of Alzheimer’s disease (AD), Dr. Alzheimer described glial lipid accumulation as one of the disease’s five pathological hallmarks^21^. Since then, glial lipid accumulation has been observed in many models–in AD mouse models, primary culture^13, 19, 22–24^, human induced pluripotent stem cell (iPSC)-derived cells^10, 12, 15, 18^, and even human brain samples^15^.

Genetics also underscore the connection between lipid metabolism and AD. Several lipid metabolism genes have been identified as modifiers of AD risk and progression^25^. Among these, coding variants of the *APOE* gene stand out as the strongest genetic risk and resilience factors for late-onset AD. The *APOE3* allele is the most common and regarded as neutral with respect to AD risk, while single missense mutations result in the *APOE4* allele (C112R), which increases AD risk, and the *APOE2* allele (R158C), which decreases AD risk^26^. Astrocytes produce the most APOE in the brain^27^ and studies in various model systems have linked *APOE4* with increased astrocytic lipid droplets^11, 12, 16, 19^.

With the rising importance of lipid droplets in Alzheimer’s disease, we aimed to systematically characterize the proteome and lipidome of lipid droplets in astrocytes harboring different *APOE* genotypes. Given differences in lipid metabolism between *APOE* mouse and human models^28^, we chose to investigate this question in human iPSC-derived astrocytes. We applied sucrose gradient ultracentrifugation to isolate lipid droplets^3^ from iPSC-derived astrocytes and subjected these lipid droplets to mass-spectrometry based proteomic and lipidomic analyses. These analyses identified, for the first time, proteins that associate with lipid droplets in human astrocytes. While many of these proteins had been found to associate with lipid droplets in other cell types, some were unique to astrocytes and suggested astrocyte-specific lipid droplet functions. Moreover, we found that *APOE* genotype influenced both the lipid and protein composition of lipid droplets, indicating that AD-associated genetic variation can reshape astrocytic cell biology via remodeling of the lipid droplet landscape. Specifically, we identified that lipid droplets in *APOE2* and *APOE4* astrocytes, which both accumulate to high levels, have different propensities for lipophagy which impact their dynamics. Together, we present a resource for the lipid droplet and neurodegenerative disease communities, offering new insight into how the most impactful genetic modifier of AD risk, *APOE,* can alter the composition and dynamics of an organelle increasingly implicated in brain health and disease.

## Results

### Sucrose gradient ultracentrifugation isolates lipid droplets from human iPSC-derived astrocytes

Astrocytes produce the majority of APOE in the brain and accumulate different numbers of lipid droplets depending on their *APOE* genotype^12, 22^. To ensure rigor and reproducibility, we chose to isolate lipid droplets from human iPSC-derived astrocytes derived from two iPSC sets (KOLF2.1J and BIONi037)^29, 30^(**Figure 1A**). Each cell set contains three cell lines that are isogenic at all loci except for *APOE*, where they are homozygous for each of the common variants: *APOE2, APOE3,* or *APOE4.* We differentiated these cells into iPSC-derived astrocytes using established protocols^31, 32^. These cells express canonical astrocytic markers (Figure S1A,B) and accumulate lipid droplets at different levels. We have previously reported findings of increased lipid droplets in *APOE4* astrocytes compared to *APOE3* astrocytes^12^. We recapitulate this finding and observe that like *APOE4, APOE2* iPSC-derived astrocytes also accumulate lipid droplets relative to *APOE3* astrocytes (**Figure 1B,C**). Given the recent research showing that lipid droplets influence disease-associated processes in AD^12, 13, 15, 18, 19, 23, 26, 33, 34^, we hypothesized that the molecular composition of the lipid droplets in *APOE2* astrocytes and *APOE4* astrocytes was different and could relate to the alleles’ divergent effects on AD risk.

**Figure 1:**
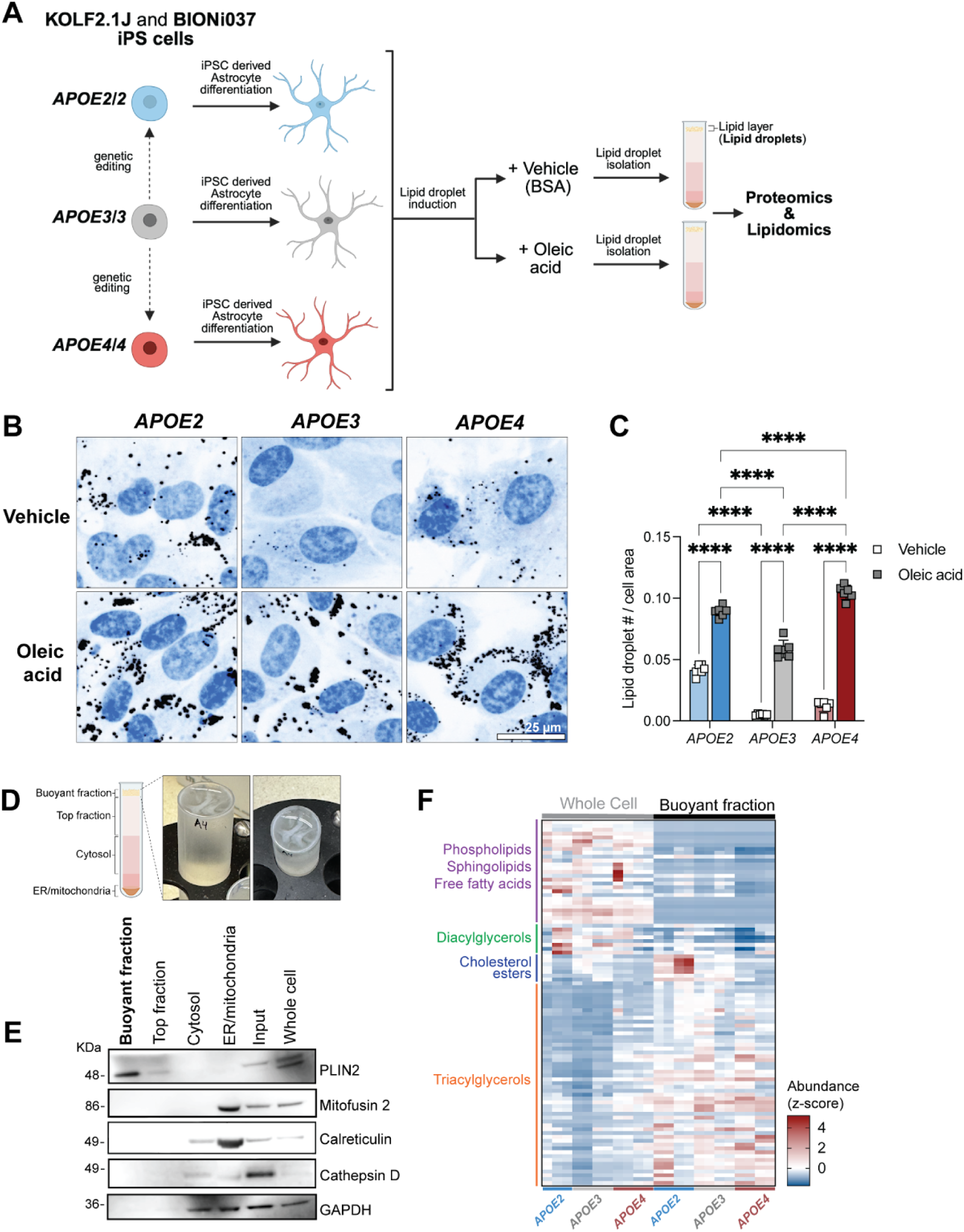
Lipid droplets can be isolated from human iPSC-derived astrocytes. A. A schematic of the lipid droplet isolation and analysis workflow from two isogenic sets of iPSC-derived astrocytes containing lines homozygous for *APOE2*, *APOE3,* and *APOE4*, from two different parental lines (KOLF2.1J and BIONi037A). B. Immunofluorescence of neutral lipids (LipidSpot, black) in isogenic iPSC-derived astrocytes (KOLF2.1J-derived) homozygous for three common *APOE* genotypes in the presence of either vehicle (BSA) or 80 μM oleic acid. Blue staining is Hoechst 33258 (nuclei). Scale bar = 25 μm. C. Quantification of lipid droplets per cell area in isogenic iPSC-derived astrocytes (KOLF2.1J-derived) homozygous for three common *APOE* genotypes in the presence of either vehicle (BSA) or 80 μM oleic acid. Each point represents an independent treatment. Data are represented as mean ± SD. **** *P* ≤ 0.0001 by two-way ANOVA with post-hoc Šídák test. D. Pictures of the buoyant fraction after sucrose gradient ultracentrifugation. E. A western blot probing the buoyant fraction for canonical markers of different cellular organelles and compartments: PLIN2 for lipid droplets, Mitofusin 2 for mitochondria, Calreticulin for endoplasmic reticulum (ER), Cathepsin D for lysosomes, and GAPDH for cytosol. F. A heatmap comparing mass spectrometry lipidomics measurements (z-score of abundance normalized to total lipid content for each sample) of un-fractionated “Whole cell” samples and buoyant fraction samples following fractionation. N=11-12 independent iPSC-derived astrocyte samples and fractionations are shown with representation from multiple lipid classes.

To maximize the number of lipid droplets in our iPSC-derived astrocytes, we stimulated lipid droplet biogenesis by treatment with oleic acid and a vehicle control (bovine serum albumin, BSA). Under these conditions, all tested iPSC-derived astrocyte lines accumulate more lipid droplets compared to vehicle control (**Figure 1B,C**,S1C,D) without substantially altering the overall lipid composition of the cell (Figure S1E). We then applied sucrose gradient density ultracentrifugation protocols to isolate lipid-rich “buoyant” fractions from iPSC-derived astrocytes treated with either oleic acid or vehicle (**Figure 1D**). This buoyant fraction was lipid droplet-rich and did not contain other contaminating organelles including lysosomes, mitochondria, and ER (**Figure 1E**). When compared to the whole cell by lipidomics, the buoyant fraction was enriched for lipids canonically present in lipid droplets like triacylglycerols and cholesterol esters (**Figure 1F**). Together, these findings confirmed that our buoyant fractions were enriched in lipid droplets and could be further analyzed to reveal molecular features of these understudied organelles in iPSC-derived astrocytes.

We first applied mass spectrometry-based proteomics to multiple cellular fractions (buoyant, cytosolic) from each astrocyte line. When comparing proteins in the oleic acid- and vehicle-treated samples, we discovered that only 26 proteins (out of 3,715 proteins detected) were differentially present in the buoyant fraction between oleic acid- and vehicle-treated conditions (Figure S2A). Of the 18 proteins specific to the buoyant fractions isolated from oleic-acid-treated iPSC-derived astrocytes, 6 were canonical lipid droplet proteins (Figure S2A), likely resulting from the greater number of lipid droplets present in the oleic acid-treated cells. Given that the amount and identity of buoyant fraction proteins detected was largely similar between oleic acid- and vehicle-treated conditions, we performed all subsequent analyses on oleic-acid treated fractions to maximize coverage of lipid droplet-associated proteins.

### The human iPSC-derived astrocyte lipid droplet-associated proteome contains known and novel proteins

Our first goal after successfully isolating lipid droplet-enriched fractions was to establish a cell type-specific lipid droplet-associated proteome for iPSC-derived astrocytes. Most past studies of lipid droplet proteomes were performed in non-human organisms or cancer cell lines^3, 6, 35^. Given the emerging role of lipid droplets in brain diseases and the central role of astrocytes in brain lipid homeostasis^22, 26^, we aimed to define a human iPSC-derived astrocyte lipid droplet-associated proteome.

We started with the 3,715 proteins detected in both technical replicates of any of our oleic acid-treated buoyant fraction samples. Of these 3,715 proteins detected, 653 were unique to the buoyant fraction and absent in the cytosolic fraction and 3,062 proteins were present in both fractions (**Figure 2A**). To determine the iPSC-derived astrocyte lipid droplet-associated proteome, we considered these two sets (unique to buoyant fraction and shared between buoyant and cytosolic fraction) separately. We first used a mean z-score > -1 filter on the 652 proteins unique to the buoyant fraction samples to select the most abundant 364 proteins (**Figure 2A,B**). These selected proteins included many canonical and well-established lipid droplet proteins such as PLIN2, PNPLA2 (ATGL), and DHRS3, which we were able to confirm localized to lipid droplets using immunofluorescence (**Figure 2C**). For the 3,062 proteins in the buoyant fraction that were also detected in cytosolic fractions, we calculated the relative enrichment of these proteins between the buoyant and cytosolic fractions and included only 76 proteins (those with the top 2.5% of enrichment scores) in our iPSC-derived astrocyte lipid droplet-associated proteome (**Figure 2A,D**). Among these 76 proteins were multiple known lipid droplet proteins that had been previously identified and validated in other studies, like ACSL3 and DHRS1^36, 37^. These proteins are known to traffic to the lipid droplet from other cellular compartments. We confirmed the lipid droplet localization of ACSL3 and DHRS1 in iPSC-derived astrocytes by immunofluorescence (**Figure 2E**).

**Figure 2:**
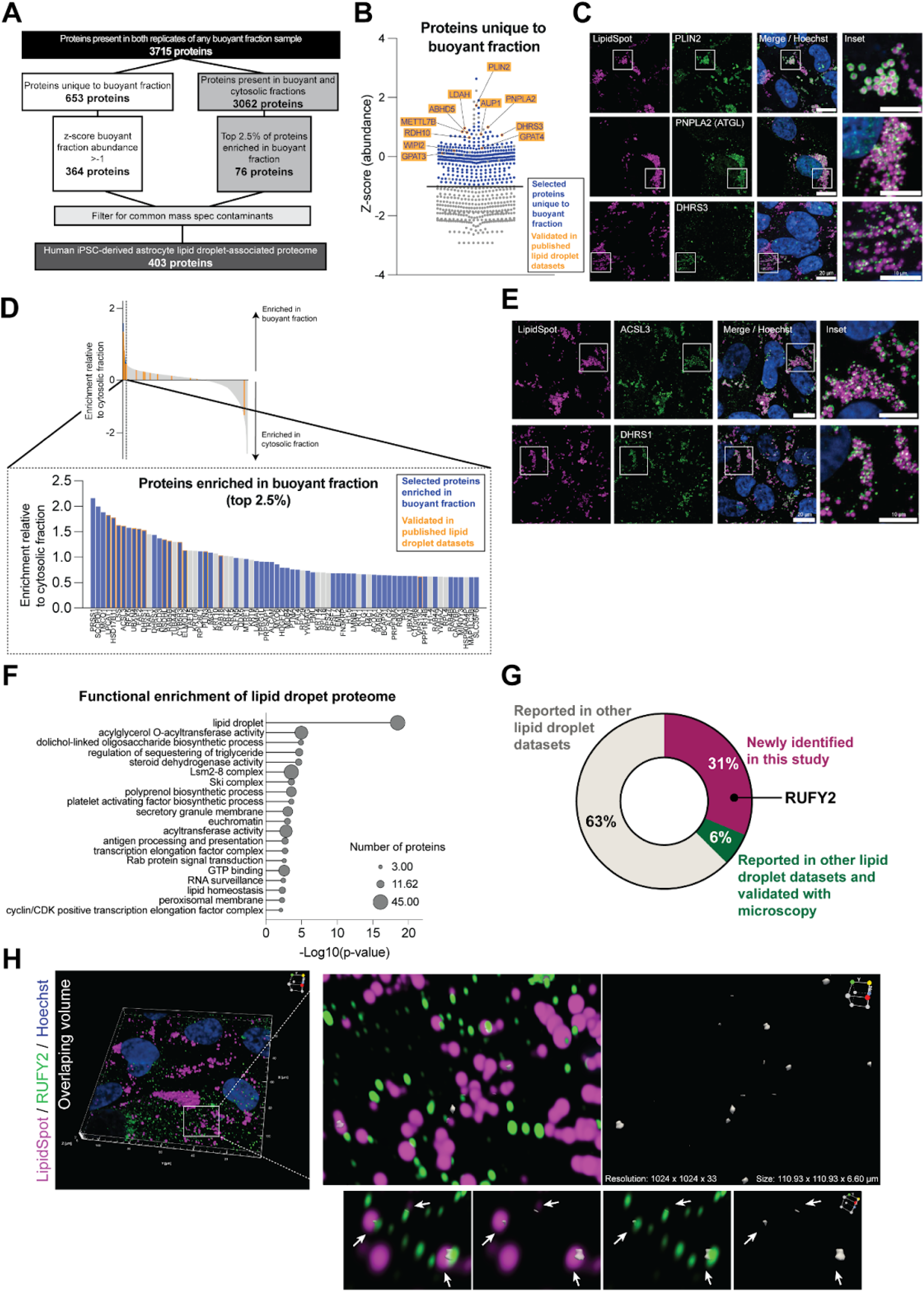
Defining the human astrocytic lipid droplet-associated proteome. A. A flowchart depicting our analysis of proteomic data to identify the human iPSC-derived astrocyte lipid droplet-associated proteome. B. A plot displaying mean z-score across all oleic-acid treated samples of any protein detected in at least both technical replicates of a buoyant fraction sample and not in the cytosolic fraction. Blue dots represent proteins selected as part of the iPSC-derived astrocyte lipid droplet-associated proteome in this study (mean z-score > -1). Orange outlines indicate canonical lipid droplet proteins that have been validated using immunofluorescence in either this or other studies. C. Immunofluorescence staining of canonical lipid droplet proteins (PLIN2, PNPLA2 (ATGL) and DHRS3) in green localized to lipid droplets (magenta) in iPSC-derived astrocytes (shown in KOLF2.1J line). Scale bar in zoomed out images is 20 μm, inset scale is 10 μm. D. Waterfall plot depicting the enrichment score relative to the cytosolic fraction of proteins present in both buoyant and cytosolic fractions. Inset depicts the top 2.5% of proteins included in our astrocytic lipid droplet-associated proteome (blue bars). Some proteins in the top 2.5% are excluded because they are common mass spectrometry experiment contaminants (grey bars). Selected proteins include multiple canonical lipid droplet proteins that have been validated using immunofluorescence in either this or other studies (orange outline). E. Immunofluorescence staining of canonical lipid droplet proteins (ACSL3 and DHRS1) in green localized to lipid droplets (magenta) in human iPSC-derived astrocytes (shown in KOLF2.1J line). Scale bar in zoomed-out images is 20 μm, inset scale is 10 μm. F. Overrepresentation analysis to determine functional enrichment of the iPSC-derived astrocyte lipid droplet-associated proteome (403 proteins). G. Pie chart depicting the percentage of lipid droplet-associated proteins uniquely identified in this study. H. Immunofluorescence staining of a novel lipid droplet-associated protein identified in this study (RUFY2) in green that contacts lipid droplets (magenta) in iPSC-derived astrocytes (shown in BIONi037 line). Arrows indicate lipid droplets interacting with RUFY2. Overlap volume is shown in white. Scale of 3D projection shown is 111 x 111 x 6.60 μm^3^.

We combined these two categories of proteins, those unique to the buoyant fraction (364) and those strongly enriched in the buoyant over the cytosolic fraction (76), to compile an initial human iPSC-derived astrocyte lipid droplet-associated proteome. Because contaminants are known to be captured in mass spectrometry isolation experiments^38^, we removed common contaminants to establish a more precise iPSC-derived astrocyte lipid droplet-associated proteome consisting of 403 proteins (Figure S2B). We used overrepresentation analysis^39^ to determine the functional enrichment of these proteins and reassuringly found that they primarily were associated with lipid droplets and lipid metabolism (**Figure 2F**). In addition to the canonical functions in lipid storage, the iPSC-derived astrocyte lipid droplet-associated proteome was enriched in functions that have not been previously associated with lipid droplets. These include functions related to RNA biology and antigen presentation. Interestingly, antigen presentation has recently been linked to astrocyte lipid state but not through the astrocytic lipid droplet proteome^32^ and major histocompatibility complex II (MHCII) molecules have been found to co-localize to lipid droplets in iPSC-derived astrocytes^40^.

Of our iPSC-derived astrocyte lipid droplet-associated proteome (403 proteins), 69% of proteins had been previously identified in other datasets. The remaining 31% of proteins were newly identified in this study (**Figure 2G**). One such protein is RUFY2, which we confirmed co-localized with lipid droplets in iPSC-derived astrocytes (**Figure 2H**), but also associates with other organelles including early endosomes and the golgi (Figure S2C). Together, our proteomics analysis revealed a high-confidence lipid droplet-associated proteome in iPSC-derived astrocytes, and identified novel proteins that distinguish glial cell lipid droplets from other cell types.

### *APOE* genotype shapes the lipid droplet-associated proteome

Given that the *APOE* genotype influences lipid droplet abundance in astrocytes, we sought to determine how it alters the lipid droplet-associated proteome. To ensure robustness, we focused exclusively on proteins and abundance patterns consistently observed across both iPSC lines sharing the same *APOE* genotype from two different isogenic sets. When comparing proteins within the lipid droplet buoyant fractions across genotypes, we found that although the majority of proteins were shared, each genotype exhibited a distinct subset of proteins (**Figure 3A**). As expected, functional enrichment analysis of the shared proteome reflected the pathways present in the iPSC-derived astrocyte lipid droplet-associated proteome (Figure S3A). The proteins exclusive to each *APOE* genotype were enriched in genotype-specific pathways. For *APOE2* these included immune pathways, for *APOE3,* metabolic signaling pathways, and for *APOE4,* protein homeostasis and autophagy pathways (**Figure 3B**). These findings suggest *APOE* genotype-specific roles for lipid droplets beyond that of lipid storage. Studies in other models have identified lipid droplets as immune hubs that dock immune proteins^4^. Lipid droplets have also been observed to participate in ER-associated protein degradation^41^.

**Figure 3:**
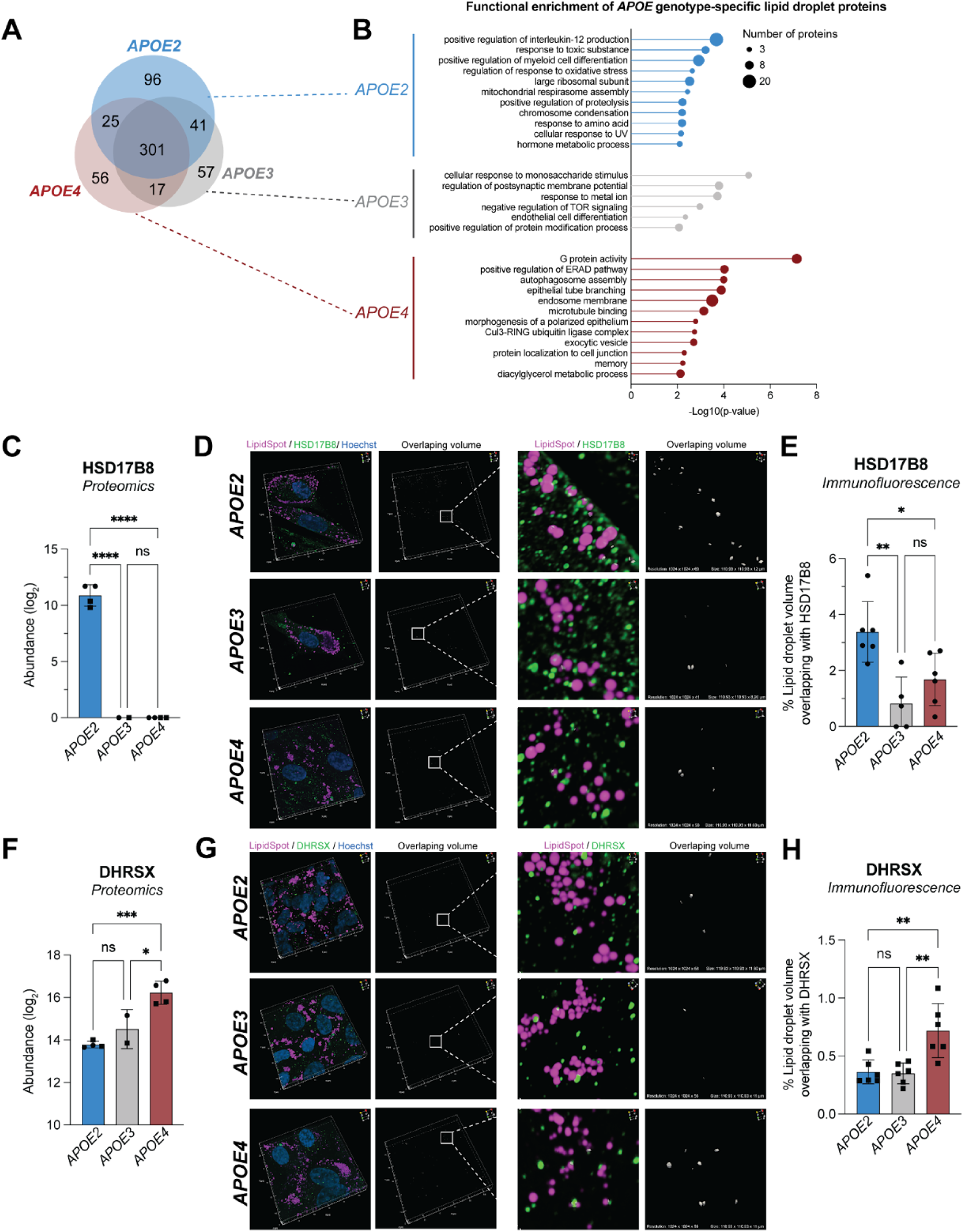
*APOE* genotype changes the lipid droplet-associated proteome. A. A Venn diagram depicting proteins in the buoyant fraction that were shared between and specific to iPSC-derived astrocytes of the three common *APOE* alleles. B. Overrepresentation analysis to determine functional enrichment of proteins exclusive to the lipid droplet-associated proteome of each *APOE* genotype. C. Abundance of HSD17B8 detected in the buoyant fractions of various *APOE* genotypes by mass spectrometry. Data represent n=4 independent replicates across two isogenic sets of human iPSC-derived astrocytes. Data are represented as mean ± SD. **** *P* ≤ 0.0001 by one-way ANOVA with post-hoc Tukey’s test. D. Immunofluorescence staining of HSD17B8 in green that contacts lipid droplets (magenta) in human BIONi037 iPSC-derived astrocytes. Overlap volume is shown in white. Scale of 3D projection shown is 111 x 111 x 12.6 μm^3^. E. Quantification of percent of lipid droplet volume overlapping with HSD17B8 signal in 3D confocal imaging. Data represent quantification of 400-700 lipid droplets per imaging frame. Data plotted are n=5-6 imaging frames per *APOE* genotype. Data are represented as mean ± SD. * *P* ≤ 0.05, ** *P* ≤ 0.01, by one-way ANOVA with post-hoc Tukey’s test. F. Abundance of DHRSX detected in the buoyant fractions of various *APOE* genotypes by mass spectrometry. Data represent n=4 independent replicates across two isogenic sets of human iPSC-derived astrocytes. Data are represented as mean ± SD. * *P* ≤ 0.05, *** *P* ≤ 0.0001 by one-way ANOVA with post-hoc Tukey’s test. G. Immunofluorescence staining of DHRSX in green that contacts lipid droplets (magenta) in human KOLF2.1J iPSC-derived astrocytes. Overlap volume is shown in white. Scale of 3D projection shown is 111 x 111 x 12.6 μm^3^. H. Quantification of percent of lipid droplet volume overlapping with DHRSX signal in 3D confocal imaging. Data represents quantification of 700-1,000 lipid droplets per imaging frame. Data plotted are n=6 imaging frames per *APOE* genotype. Data are represented as mean ± SD. ** *P* ≤ 0.01 by one-way ANOVA with post-hoc Tukey’s test.

We also identified proteins that were associated with lipid droplets in all three *APOE* genotypes but to different extents among the genotypes (Figure S3B). For example, we observed that HSD17B8, a protein involved in steroid and fatty acid metabolism^42^, was only detected in our *APOE2* astrocytes (**Figure 3C**). Using immunofluorescence we observed that HSD17B8 showed greater colocalization with lipid droplets in *APOE2* astrocytes than in *APOE3* or *APOE4* astrocytes (**Figure 3D,E**). DHRSX, a protein involved in multiple steps of dolichol biosynthesis^43^, was more abundantly associated with *APOE4* lipid droplets than in the other two *APOE* genotypes (**Figure 3F**) and we were able to confirm this with immunofluorescence (**Figure 3G,H**). Together, these findings cement that iPSC-derived astrocytes harboring different *APOE* genotypes have distinct lipid droplet-associated proteomes. These genotype-specific proteins may underlie functional differences in lipid droplet biology between astrocytes.

### *APOE* genotype shapes the lipid droplet lipidome

We hypothesized that the effect of *APOE* genotype on the astrocytic lipid droplet-associated proteome may arise from changes to lipid droplet lipid composition. To investigate this, we analyzed the lipid composition of lipid droplets from oleic acid- and vehicle-treated iPSC-derived astrocytes of various *APOE* genotypes using LC-MS-based lipidomics. When we compared the overall lipid profile of the buoyant fraction with that of the whole cell and the ER/mitochondria fraction, we observed that each cellular compartment had a unique lipid profile (**Figure 4A**). In addition, we noticed that some *APOE* genotype-dependent lipid changes at the organellar level were not captured in the whole cell data. This finding reveals that *APOE* genotype modulates lipid composition at the subcellular level, and highlights the importance of studying lipid biology from an organellar perspective.

**Figure 4:**
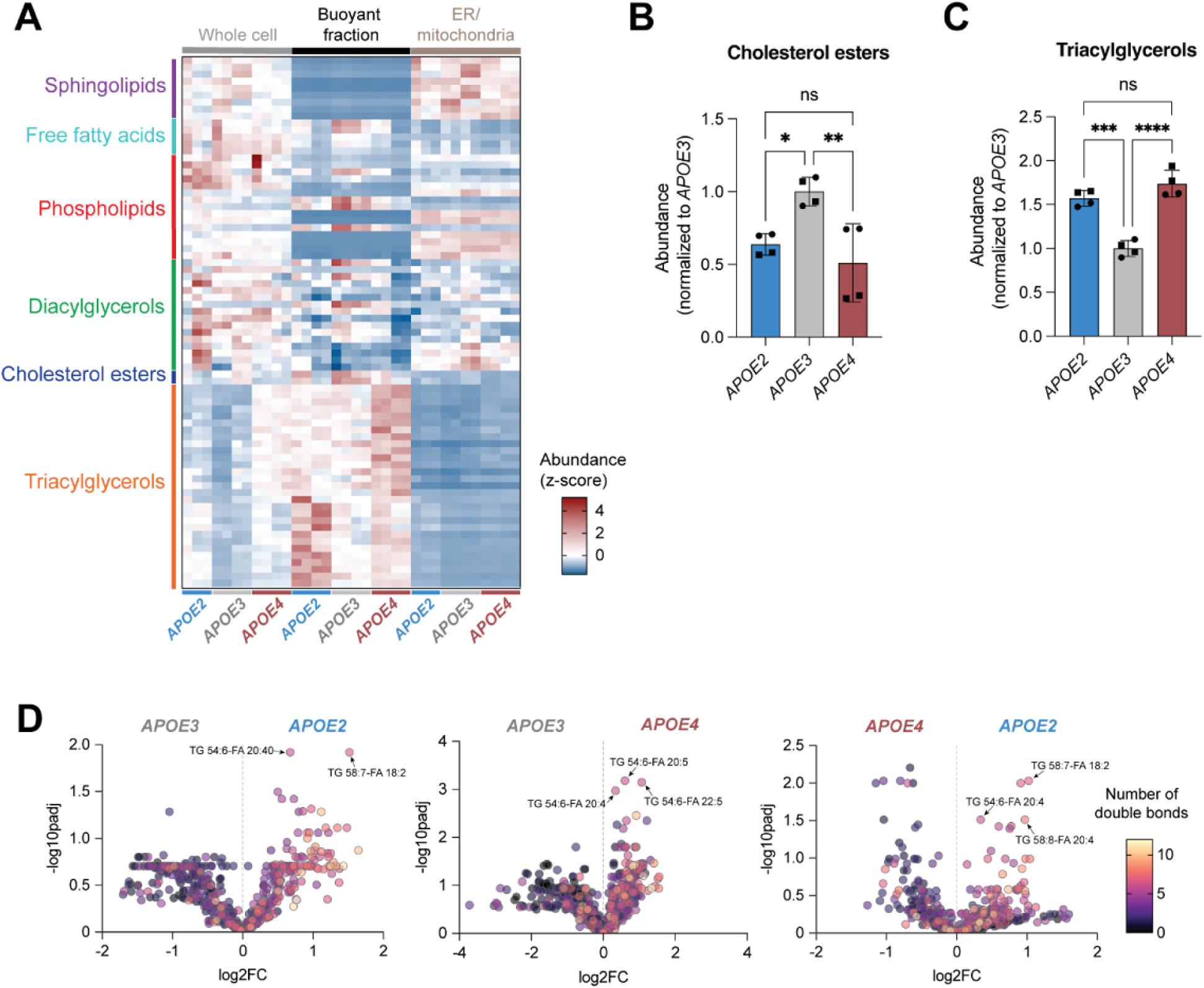
*APOE* genotype changes the lipid droplet lipidome. A. Heatmap depicting mass spectrometry lipidomics measurements (z-score of abundance normalized to total lipid content) of un-fractionated whole cells, as well as buoyant (lipid droplet) and ER/mitochondria fractions. Samples were obtained from iPSC-derived astrocytes treated with oleic acid. N=11-12 independent iPSC-derived astrocyte samples and fractionations are shown with representation from multiple lipid classes. B. Abundance of cholesterol esters in buoyant fractions of various *APOE* genotypes obtained from iPSC-derived astrocytes treated with oleic acid. Data represent n=4 independent replicates across two isogenic sets of human iPSC-derived astrocytes. * *P* ≤ 0.05, ** *P* ≤ 0.01 by one-way ANOVA with post-hoc Tukey’s test. C. Abundance of triacylglycerols in buoyant fractions of various *APOE* genotypes obtained from iPSC-derived astrocytes treated with oleic acid. Data represent n=4 independent replicates across two isogenic sets of human iPSC-derived astrocytes. *** *P* ≤ 0.001, **** *P* ≤ 0.0001 by one-way ANOVA with post-hoc Tukey’s test. D. Volcano plots comparing buoyant fraction lipids across different *APOE* genotypes obtained from iPSC-derived astrocytes treated with oleic acid. Color coding represents fatty acid saturation (number of double bonds).

When comparing the levels of canonical lipid droplet lipids (cholesterol esters and triglycerides) and their precursors (diacylglycerols and fatty acids) after oleic acid treatment in the buoyant fraction, we observed that these correlated well to the quantity of lipid droplets in the iPSC-derived astrocytes of various *APOE* genotypes (**Figure 4B,C**, S4A,B). Lipid droplets from *APOE3* iPSC-derived astrocytes showed high cholesterol esters and fatty acids but reduced triacylglycerols, suggesting a lower capacity to efficiently store fatty acids as neutral lipids when stimulated with oleic acid (**Figure 4B,C**, S4B). On the other hand, lipid droplets from *APOE2* and *APOE4* iPSC-derived astrocytes had similar quantities of cholesterol esters and triacylglycerols (**Figure 4B,C**). Because the lipid droplet-associated proteomes of these genotypes were distinct, we reasoned that changes in relative lipid class were not the driver of lipid droplet-associated proteome changes.

Previous work has identified that *APOE* genotype correlates with changes in fatty acid saturation in whole cell lipidomics measurements^11, 12, 32^. When examining the saturation of fatty acids within lipid droplet lipids, we observed that *APOE2* contains far more unsaturated lipids than *APOE3* and *APOE4* (**Figure 4D**). This change was not reflected in our whole cell lipidomics data (Figure S4C) reinforcing the importance of cellular compartment-specific analyses. Given that our experimental paradigm involved treating our iPSC-derived astrocytes with oleic acid, a monounsaturated fatty acid, to increased lipid droplet content, we confirmed that the trend we observed of increased unsaturation in *APOE2* lipid droplets held true under basal conditions (treated with BSA vehicle) (Figure S4D). Differences in saturation between genotypes were mainly driven by triacylglycerol containing polyunsaturated fatty acid (PUFA) species, such as TG 54:6-FA 20:4 and TG 58:7-FA 18:2 (**Figure 4**, S4D). Several biophysical studies have reported that lipid droplets with higher levels of unsaturation exhibit packing defects and increased protein binding^44, 45^. Moreover, the sequestration of membrane PUFA lipids into lipid droplets could be protective, as it prevents peroxidation and subsequent lipotoxicity^46^. Given that lipid droplets from *APOE2* astrocytes had both the highest unsaturation of lipids and the greatest number of proteins bound, we concluded that *APOE* genotype could be driving changes in lipid droplet-associated proteomes by modulating lipid saturation.

### Lipid droplet-associated proteomes modulate lipid droplet dynamics

Since *APOE* genotype modulates both the lipid droplet-associated proteome and lipidome in iPSC-derived astrocytes, we asked whether these changes have functional consequences. One of the enriched functional categories in the *APOE4* lipid droplet-associated proteome was autophagy (**Figure 3B**). Our dataset revealed that in *APOE4* astrocytes, lipid droplets contained higher levels of autophagy-related proteins than *APOE2* astrocytes, even though both *APOE2* and *APOE4* astrocytes exhibit higher numbers of lipid droplets than *APOE3* astrocytes. One interesting candidate is SQSTM1 (p62). SQSTM1/p62 is an adaptor protein that binds ubiquitinated proteins and organelles and recruits them to autophagosomes for degradation^47^. Our proteomics data revealed that SQSTM1/p62 was more abundant on *APOE4* lipid droplets than *APOE2* lipid droplets (**Figure 5A**). We confirmed this by immunofluorescence and observed higher levels of SQSTM1/p62 co-localized with lipid droplets in *APOE4* than *APOE2* astrocytes (**Figure 5B,C**, S5A,B).

**Figure 5:**
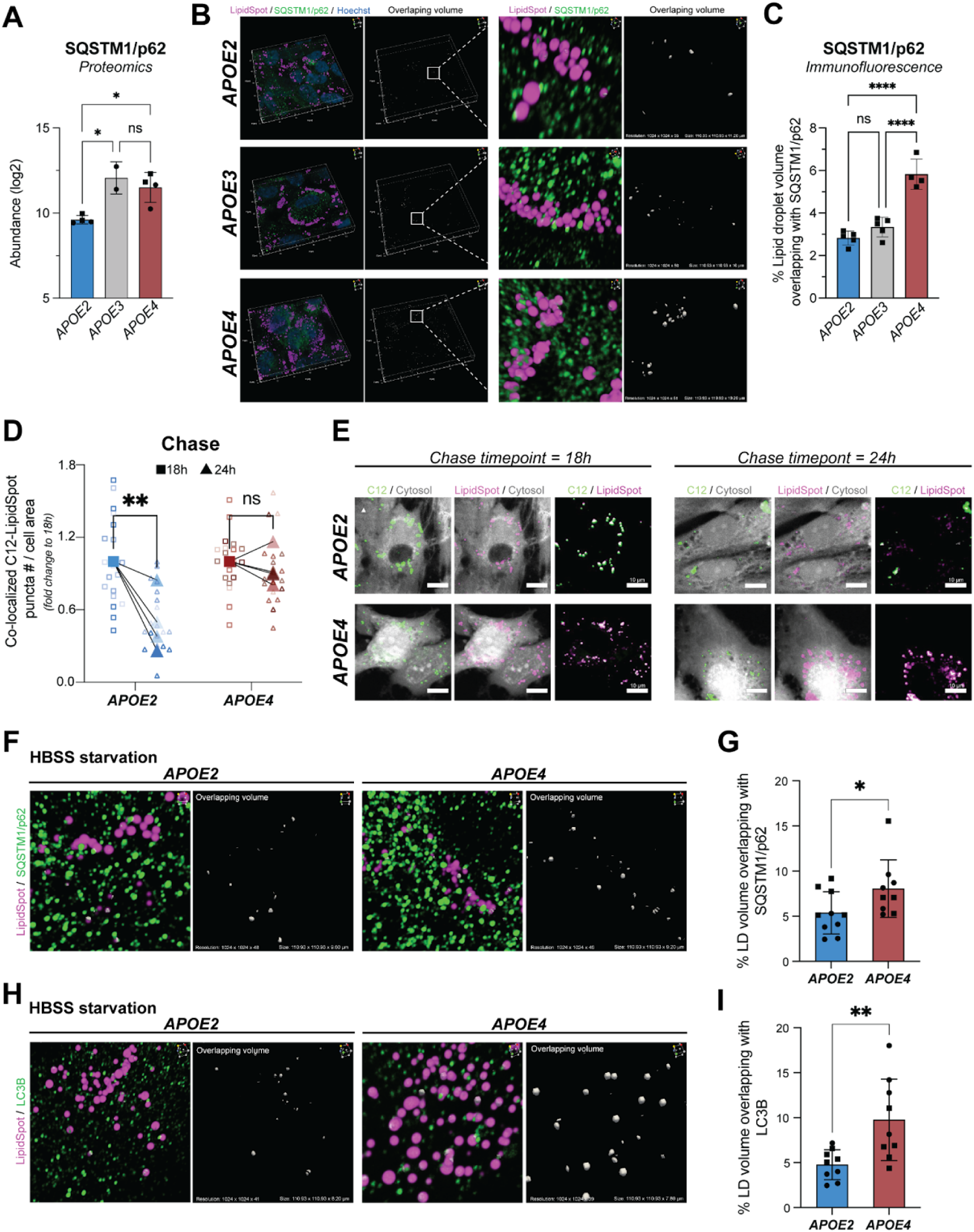
*APOE* genotype-specific lipid droplet-associated proteins change lipophagy. A. Abundance of SQSTM1/p62 detected in the buoyant fractions of various *APOE* genotypes by mass spectrometry. Data represent n=4 independent replicates across two isogenic sets of human iPSC-derived astrocytes. Data are represented as mean ± SD. * *P* ≤ 0.05 by one-way ANOVA with post-hoc Tukey’s test. B. Immunofluorescence staining of SQSTM1/p62 in green that contacts lipid droplets (magenta) in human iPSC-derived astrocytes harboring the three *APOE* genotypes (shown in KOLF2.1J line). Overlap volume is shown in white. Scale of 3D projection shown is 110.93 x 110.93 x 10-11.20 μm^3^. C. Quantification of percent of lipid droplet volume overlapping with SQSTM1 signal in 3D confocal imaging (shown in KOLF2.1J line). Data represent quantification of an average of ∼700 lipid droplets per imaging frame. Data plotted are n=4-5 imaging frames per *APOE* genotype. Data are represented as mean ± SD. **** *P* ≤ 0.0001 by one-way ANOVA with post-hoc Tukey’s test. D. The change in number of BODIPY-C_12_-positive lipid droplets (co-labeled with LipidSpot) during a chase period of 18 h and 24 h in human iPSC-derived astrocytes harboring *APOE2* and *APOE4* genotypes (shown in KOLF2.1J line). Data shown in large symbols represent n=4 independent pulse chase experiments calculated as fold change to 18 h. Data shown in small symbols represent individual replicates for each experiment. ** *P* ≤ 0.01 by repeated measures two-way ANOVA with post-hoc Šidák’s test. E. Immunofluorescence depicting co-localized BODIPY-C_12_-positive lipid droplets at chase timepoints 18 h and 24 h (shown in KOLF2.1J line). Scale bars are 10 μm. F. Immunofluorescence staining of SQSTM1/p62 in green that contacts lipid droplets (magenta) in *APOE2* and *APOE4* iPSC-derived astrocytes under HBSS-induced starvation (shown in BIONi037 line). Overlap volume is shown in white. Scale of 3D projection shown is 110.93 x 110.93 x 9.20-9.61 μm^3^. G. Quantification of percent number of lipid droplet volume co-localizing with SQSTM1/p62 following starvation by HBSS for 1 h shown across two isogenic sets of human iPSC-derived astrocytes. Data represent quantification of an average of ∼100 lipid droplets per condition. Data plotted are n=9-10 imaging frames per *APOE* genotype. Data are represented as mean ± SD. *P* = 0.0502 by unpaired t-test. H. Immunofluorescence staining of LC3B in green that contacts lipid droplets (magenta) in *APOE2* and *APOE4* iPSC-derived astrocytes under HBSS-induced starvation (shown in BIONi037 line). Overlap volume is shown in white. Scale of 3D projection shown is 110.93 x 110.93 x 7.80-8.20 μm^3^. I. Quantification of percent number of lipid droplet volume co-localizing with LC3B following starvation by HBSS for 1 h shown across two isogenic sets of human iPSC-derived astrocytes. Data represent quantification of an average of ∼100 lipid droplets per condition. Data plotted are n=9 imaging frames per *APOE* genotype. Data are represented as mean ± SD. ** *P* ≤ 0.01 by unpaired t-test.

One route for lipid droplet turnover is via lipophagy–the autophagy of lipid droplets–which involves ubiquitination and binding to autophagy adaptor proteins like SQSTM1/p62^1^. Given the differences in lipid droplet-associated proteome composition, we hypothesized that *APOE4* astrocytes may display an impairment in lipophagy leading to an accumulation of lipid droplets bound to SQSTM1/p62^48^. To test this hypothesis, we first measured the overall turnover rate of fatty acids within lipid droplets in *APOE2* and *APOE4* astrocytes by using an established pulse-chase experimental paradigm with fluorescent BODIPY-C_12_ fatty acids^49^. C_12_ fatty acids within lipid droplets persisted longer in *APOE4* than in *APOE2* astrocytes following a 24h chase period (**Figure 5D,E**, S5C). To address whether *APOE* genotype differentially impacts lipophagy under conditions where we induced autophagy, we analyzed the co-localization of lipid droplets and SQSTM1/p62 or the autophagosome marker LC3B following acute HBSS-induced starvation. We observed that *APOE4* lipid droplets showed greater co-localization with both SQSTM1/p62 and LC3B than *APOE2* lipid droplets (**Figure 5F,G**). This finding suggests that lipophagy in *APOE4* astrocytes is impaired and cannot be activated by starvation-induced autophagy. Given that *APOE2* and *APOE4* astrocytes both accumulate lipid droplets under basal conditions, we hypothesized that prolonged starvation would decrease the number of *APOE2* lipid droplets but not impact *APOE4* lipid droplets (as they are resistant to lipophagy). We observed that *APOE2* astrocytes exhibited a trend towards reduced lipid droplets while *APOE4* astrocytes did not (Figure S5D). Together these findings suggest that although both genotypes accumulate significant numbers of lipid droplets, *APOE4* lipid droplets are resistant to lipophagy while *APOE2* lipid droplets are not. Taken together, these findings demonstrate that characterizing the *APOE* genotype-specific lipid droplet-associated proteome provides a mechanistic basis for understanding genotype-specific lipid droplet dynamics.

## Discussion

Lipid droplets are dynamic organelles that impact neurodegenerative disease biology. Lipid droplet proteins are key to specifying their many functions. For the first time, we defined the protein and lipids associated with lipid droplets isolated from human iPSC derived astrocytes homozygous for the *APOE2* (protective), *APOE3* (neutral), and *APOE4 (*risk) genotypes.

Our analyses reveal that human astrocytic lipid droplets exhibit molecular features shared with many cell types and others that are specific to astrocytes. We confirmed the presence of canonical lipid droplet proteins (PLIN2, PNPLA2, ACSL3, DHRS1) and observed that astrocytic lipid droplet-associated proteins have non-canonical functions like antigen presentation. We identified novel lipid droplet-associated proteins like RUFY2, a protein involved in endosomal trafficking. RUFY2 contains a PI3P-binding FYVE domain, linking it to lipid droplet metabolism, a connection previously established for other FYVE-containing proteins like DFCP1^50, 51^. Genetic knockdown studies show RUFY2 influences lipid droplet morphology and distribution^9, 35^, hinting at a potential link between vesicle trafficking defects^52^ and altered lipid biology in astrocytes, particularly in neurodegenerative contexts.

When examining the effects of *APOE* genotype on the composition of lipid droplets, we discovered that the accumulation of lipid droplets is not inherently pathological; rather, their impaired dynamics and turnover distinguishes disease risk from protection. Although both *APOE4* and *APOE2* astrocytes accumulate more lipid droplets than *APOE3*, the molecular features of lipid droplets are different. *APOE2*-associated droplets harbor immune proteins, pointing to potential immunomodulatory roles that may underlie protective biology. This finding mirrors observations in other systems where lipid droplets regulate immune responses^4, 53^. *APOE4* lipid droplets accumulate stalled lipophagy adapter proteins and display impaired lipophagy compared to *APOE2* lipid droplets, suggesting that the autophagy-mediated turnover of lipid droplets differentiates risk and protective *APOE* genotypes. Our findings align with a recent study on human data that identified lipophagy genes as dysregulated in AD patients and AD risk^54^. Previous work has linked disruptions to autophagy and lysosomal biology with Alzheimer’s disease^55–57^. Studies in other model systems have observed impaired autophagy in *APOE4* astrocytes and other glia^11, 58, 59^. Similarly, reduced lipophagy has been shown to contribute to the progression of AD in disease mouse models^60^. Together with our results, these findings reveal that *APOE* genotype alters lipophagy via changes to the lipid droplet-associated proteome and that lipophagy disruptions could contribute to *APOE4* associated risk.

*APOE* genotype also impacts lipid saturation within lipid droplets. *APOE2* lipid droplets exhibit higher levels of PUFA-enriched triacylglycerols compared to *APOE4* and *APOE3* lipid droplets. While PUFAs in membrane phospholipids can promote ferroptosis through lipid peroxidation, their sequestration in the triacylglycerol-rich lipid droplet cores mitigates ferroptosis and lipotoxicity^46^. Additionally, PUFAs within lipid droplets can be metabolized into lipid signalling mediators in the inflammatory response^46^. We speculate that *APOE2*-mediated protection in astrocytes could arise from their ability to dynamically mobilize lipid droplet lipids to respond to inflammatory stressors implicated in neurodegenerative diseases. The less labile *APOE4* lipid droplets are less able to respond and buffer cellular stress and therefore present a liability to cell health when faced with disease-associated stresses.

There are many natural extensions of this work for future studies. We have uncovered genotype and cell type-driven diversity in lipid droplet composition. However, our study could not capture heterogeneity within lipid droplets in a single cell type or genotype, which may provide greater insights into organellar function^61, 62^. Our study focuses on the cell autonomous regulation of lipid droplets. Various studies have shown that lipid droplets influence intercellular communication^15, 18, 49^. In future work, we hope to understand the cell non-autonomous consequences of lipid droplet turnover and in this context, explore whether modulating lipid droplet plasticity can modulate disease risk.

Given the centrality of lipid and glial cell dysregulations contributing to neurodegenerative disease, our study sheds light on the diverse and understudied roles of lipid droplets in human astrocytes. Moreover, understanding *APOE* genotype-driven differences can illuminate the molecular mechanisms underlying the biology of genetic risk and protection for AD.

## Methods

### Human iPSC lines and culture

Isogenic BIONi037-A (APOE3/3), BIONi037-A-2 (APOE2/2) and BIONi037-A-4 (APOE4/4) human iPSC lines (77 year-old female donor) were obtained from the European Bank for induced pluripotent Stem Cells (EBiSC). Isogenic KOLF2.1J *APOE* derivatives JIPSC001000 (APOE3/3), JIPSC001154 (APOE2/2), and JIPSC001150 (APOE4/4) were obtained from The Jackson Lab/iPSC Neurodegenerative Disease Initiative human iPSC collection (57 year-old male donor). All iPSCs were maintained in either mTeSR Plus (STEMCELL Technologies 100-0276, KOLF2.1J) or Essential 8 Medium (E8; Thermo Fisher Scientific A1517001, BIONi037-A) and plated and expanded on either hESC-Qualified Matrigel Matrix (Corning 354277, KOLF2.1J) or GelTrex (Thermo Fisher Scientific, A1413302, BIONi037-A). Cells were passaged at 90% confluency with either ReLeSR (STEMCELL Technologies 100-0483, KOLF2.1J) or 1 mM EDTA (Thermo Fisher Scientific 15575-038, BIONi037-A) in 1X PBS (VWR, 392-0434). Genomic stability of all iPSC lines was characterized by karyotyping or SNP array testing and all samples were tested to ensure no mycoplasma contamination.

### iPSC differentiation to iPSC-derived astrocytes

Astrocytes were differentiated from iPSCs using small-molecule protocols. For KOLF2.1J-derived astrocytes, neural progenitor cells (NPCs) were generated using STEMdiff SMADi Neural Induction Kit (STEMCELL Technologies 08581) using the kit monolayer protocol. NPCs were then differentiated into astrocytes using human recombinant BMP-4 (Thermo Fisher Scientific 120-05ET) and human recombinant FGF-2 (Thermo Fisher Scientific 100-18B) as previously described^31^. Cells were sorted twice using ACSA-1 APC-conjugated antibodies (Miltenyi Biotec 130-123-555) and frozen at 1 x 10^6^ cells/mL in CryoStor CS10 (STEMCELL Technologies 07930) in cryovials (Azenta Life Sciences 68-1001-11). BIONi037-A iPSC lines were differentiated into astrocytes using previously described protocols^32^ Maintenance of all iPSC-derived astrocytes was performed in complete Astrocyte Media (AM; ScienCell Research Laboratories 1801) and cells were passaged at 80-90% confluence using TrypLE Express (Thermo Fisher Scientific 12604039). iPSC-derived astrocytes were cultured in AM media without fetal bovine serum (FBS; ScienCell Research Laboratories 0010) for 4 days before experiments.

### Lipid droplet isolation

iPSC-derived astrocytes were grown in 4 x 150 mm dishes (Corning 353025) until confluent and treated with oleic acid (80 μM, 48h; MilliporeSigma O3008) or bovine serum albumin (BSA, 0.7%, 48h; MilliporeSigma A3311) in AM media without FBS. Lipid droplets were isolated by density gradient centrifugation using an adapted protocol^63, 64^. Briefly, cells were washed and scraped using a Cell Lifter (Corning 3008) in ice-cold 1X DPBS (Quality Biological 114-057-101), and incubated for 10 min on ice in ice-cold hypotonic lysis medium (HLM;20 mM Tris-HCl, pH 7.4 and 1 mM EDTA) containing 1X Halt Protease and Phosphatase Inhibitor Cocktail (Thermo Fisher Scientific 78440). Cell suspensions were Dounce homogenized (Wheaton 357542) and lysates were centrifuged at 1,000 x g for 10 min (Eppendorf 5424R). The supernatant was diluted to a final concentration of 20% sucrose (MilliporeSigma S0389) in HLM and transferred to the bottom of a 14 mL Ultra-Clear round bottom tube (Beckman Coulter Life Sciences 344060). To create a sucrose gradient, 5 mL of 5% sucrose in HLM were gently layered on top, followed by 4.5-6.5 mL of HLM. Overlaid sucrose gradient samples were centrifuged at 28,000 x g for 1 h at 4°C using an SW-40 Ti swinging bucket rotor (Beckman Coulter Life Sciences 331301) in an ultracentrifuge. Buoyant fractions, which contain isolated lipid droplets, were pipetted from the top 200-500 μL of the resultant gradient. The remaining fractions (*i.e.*, cytosolic and pellet) were also collected. All fractions were stored at -80°C for immunoblotting and mass spectrometry-based proteomics and lipidomics analyses.

### Immunoblotting

The purity of buoyant fractions was analyzed by immunoblotting. Whole cell samples were lysed with 1X RIPA Lysis and Extraction buffer (MilliporeSigma 20-188) and protein concentration was estimated using the bicinchoninic acid (BCA) protein assay (Thermo Fisher Scientific 23227). For SDS-PAGE separation, whole cell protein lysates, and the buoyant, cytosolic and pellet fractions were denatured with 1X NuPAGE LDS Sample Buffer (Thermo Fisher Scientific NP0007) and 1X NuPAGE Sample Reducing Agent (Thermo Fisher Scientific NP0009), separated on NuPAGE 4-12% Bis-Tris gels (Thermo Fisher Scientific NP0323BOX) and transferred onto iBlot2 nitrocellulose Transfer Stacks (Thermo Fisher Scientific IB23002) using the iBlot2 Dry Blotting System (Thermo Fisher Scientific IB21001). Membranes were blocked with Intercept TBS blocking buffer (LICORbio 927-60001) for 1 h at room temperature, and probed overnight with primary antibodies at 4°C. After washing with 1X TBS (KD Medical RGF-3385) containing 0.1% (v/v) Tween 20 (MilliporeSigma P9416), membranes were incubated with horseradish peroxidase (HRP)-conjugated secondary antibodies (1:10,000) for 1 h at room temperature. Immunoblots were developed with SuperSignal West Pico Chemiluminescent Substrate (Thermo Fisher Scientific 34580) and imaged on an Amersham Imager 680 (Cytiva).

### Proteomics sample preparation

Sample preparation for proteomics was conducted on a fully automated workflow as previously described^65^. Briefly, 200 μL lysis buffer (50 mM Tris-HCI (Thermo Fisher Scientific 15568025), 50 mM NaCl (ThermoFisher 24740011), 1% SDS (Thermo Fisher Scientific 15553027), 1% Triton X-100 (MilliporeSigma T8787), 1% NP-40 (Thermo Fisher Scientific 85124), 1% Tween-20 (MilliporeSigma P9416), 1% glycerol (MP Biomedicals 800687), 1% sodium deoxycholate (wt/vol; MilliporeSigma D6750), 5 mM EDTA (Thermo Fisher Scientific 15575020), 5mM dithiothreitol (DTT; Thermo Fisher Scientific 20290), 5KU benzonase (MilliporeSigma E8263), and 1× complete protease inhibitor (MilliporeSigma 5892970001)) was added to the samples. The lysates were incubated at 65°C for 30 min at 1200 rpm for protein denaturation. Samples were then alkylated with 10 mM iodoacetamide (Thermo Fisher Scientific A39271) for 30 min at room temperature. The protein concentration was determined using DC Protein Assay (Bio-Rad 5000111). The protein enrichment and on-bead (MilliporeSigma GE45152105050250, GE665152105050250) tryptic/Lys-C (Promega X100) digestion were conducted using a KingFisher APEX robotic system. Proteins were digested with Trypsin/Lys-C mix (Promega V5073) in 50 mM ammonium bicarbonate (MilliporeSigma A6141) at 37°C for 18h. Following digestion, peptides were vacuum-dried and reconstituted in 2% acetonitrile (ACN, Thermo Fisher Scientific TS-51101) with 0.1% formic acid (Thermo Scientific 28905). 1μg of peptides was used for LC-MS/MS proteomic analysis.

### Liquid chromatography and mass spectrometry

For each sample, digested peptides were analyzed using an UltiMate 3000 nano-HPLC system (Thermo Fisher Scientific) coupled with an Orbitrap Eclipse mass spectrometer (Thermo Fisher Scientific). The peptides were loaded at 5 μL/min onto a trap column (75 μm × 2 cm, PepMap nanoViper C18 column, 3μm, 100 Å, Thermo Fisher Scientific) equilibrated in 2% ACN with 0.1% trifluoroacetic acid (TFA, Thermo Scientific 85183). The samples were loaded to trap column for 5 min and switched in-line with ES903A nano C18 column (75 μm × 500 mm, 2 μm, 100 Å,Thermo Scientific) using an 80-minute linear gradient of 2% - 40% phase B (5% DMSO in 0.1% formic acid, in ACN). The column temperature was maintained at 60°C and the flow rate was set to 300 nL/min. Data were acquired in data-independent acquisition (DIA) mode. The MS1 scan was set at a resolution of 120,000, with a standard AGC target, and the maximum injection time was set to auto. The MS2 scans covered a precursor mass range of 400–1000 m/z, using an isolation window of 8 m/z with 1 m/z overlap, resulting in a total of 75 windows per scan cycle. Fragmentation was performed using high-energy collisional dissociation (HCD) with a normalized collision energy of 30%. MS2 spectra were acquired at a resolution of 30,000, with a scan range of 145–1450 m/z, and the loop control was set to 3 seconds.

### Proteomics data analysis

We performed DIA database searches using Spectronaut (version 19, Biognosys) with the directDIA search strategy. We used the UniProt human proteome reference with reviewed genes (20,384 entries) as the library FASTA file. directDIA false discovery rate set to 1%, protein N-terminal acetylation and methionine oxidation were set as variable modifications and carbamidomethylation of cysteine residues was selected as a fixed modification.

### Bioinformatic analysis of the iPSC-derived astrocyte lipid droplet-associated proteome

Each fraction (cytosolic and buoyant) the Spectronaut output was subjected to filtering, normalization, and imputation using the DEP2 package in R^66^. Proteins present in each replicate of any genotype-treatment group across both lines were retained, then values were normalized using variance stabilizing normalization^67^. Imputation was carried out using a mixed imputation model where imputation for proteins with randomly missing values were imputed using kNN, and proteins not missing at random, defined as proteins missing in every replicate of a genotype-treatment group, were imputed as 1.

To determine differential abundance statistical analysis was conducted using the DEP2 package^66^ which utilizes limma and fdrtools. To identify proteins differentially abundant between oleic acid and BSA loading control treated cells, comparisons of all oleic acid samples to all BSA samples from the buoyant fraction across both lines were assessed, proteins were identified as differentially abundant if they met a p-value cutoff of <0.1. To determine proteins that were differentially abundant between genotypes buoyant fraction oleic acid samples were compared, with proteins identified as differentially abundant with an adjusted p-value threshold of <0.1.

To identify proteins that were enriched in the lipid droplet fraction compared to the cytosolic fraction. Z-score transformations were applied to the cytosolic and buoyant fraction datasets, and a mean z-score of oleic acid treated samples was calculated for each protein identified in the fractions. A linear model was fitted to the mean z-scores of the buoyant and cytosolic fractions. Only complete cases were considered. Residuals were calculated for each protein detected in the buoyant fraction. Residual scores were then ranked to determine proteins enriched in the buoyant fraction. This was repeated to identify proteins enriched in a single genotype by calculating a mean z-score, and ranked residual value considering samples from each genotype separately. All proteome lists were filtered for common mass spectrometry contaminants using a published database^38^. Proteins that appeared in over half of the mass spectrometry studies in the database were excluded from our analyses.

### Lipidomics sample preparation and analysis

Comprehensive targeted lipidomics was accomplished using a flow-injection assay based on lipid class separation by differential mobility spectroscopy and selective multiple reaction monitoring (MRM) targeting individual lipid species (Lipidyzer platform). A detailed description of lipid extraction, employed software, and the quantitative nature of the approach can be found elsewhere^68–70^. Briefly, after the addition of >60 deuterated internal standards, lipids were extracted using methyl tert-butyl ether. Organic extracts were combined and subsequently dried under a gentle stream of nitrogen and reconstituted in running buffer. Lipids were then analyzed using flow-injection in MRM mode employing a Shimadzu Nexera series HPLC and a Sciex QTrap 6500+ mass spectrometer. For further analysis, lipidomic data normalized to cell number or sample volume were used. Specifically, 25 μL of lipid droplet isolations or cell pellets were used. SODA light software was used for data visualization^32^.

### Immunofluorescence microscopy and analysis

To quantify the number of lipid droplets in iPSC-derived astrocytes under lipid droplet formation conditions, cells were plated on 96-well μ-Plates (ibidi 89626) and cultured with oleic acid (80 μM) or vehicle control (0.7% BSA) diluted in AM media without FBS. After 48h, cells were rinsed with DPBS and fixed with 4% paraformaldehyde (PFA; Electron Microscopy Sciences 15710-S). Lipid droplets were labeled with LipidSpot 610 stain (1:1,000; Biotium 70069) as per manufacturer’s instructions. Cell membranes and nuclei were counterstained with MemBrite Fix 568/580 cell surface stain (1:5,000; Biotium 30095) and Hoechst 33258 (1:10,000; Thermo Fisher Scientific H3570), respectively. Images were acquired using a Nikon CSU-W1 Spinning Disk microscope with a 40X water immersion objective lens (NA 1.25). Z-stacks with a stack height of 0.5 μm were acquired and images were analyzed using the Nikon NIS-Elements General Analysis (GA3) suite.

To determine lipid droplet co-localization of canonical and novel lipid droplet-associated proteins, iPSC-derived astrocytes were treated with oleic acid (80 μM, 48h) in AM media without FBS and fixed with 4% PFA. After fixation, cells were permeabilized with 0.01% saponin (MilliporeSigma SAE0073) in DPBS for 10 min, or with 0.3% Triton X-100 in DPBS for 5 min. Following permeabilization, cells were blocked for 1 h at room temperature with 5% BSA, 1% normal donkey serum (MilliporeSigma S30-M), 0.01% saponin in DPBS (for samples permeabilized with saponin) or 5% BSA, 1% normal donkey serum in DPBS (for samples permeabilized with Triton X-100). Primary antibodies diluted in the respective blocking buffers were incubated overnight at 4°C. Cells were washed with DPBS and secondary antibodies (1:2,000) were diluted in the respective blocking buffers and incubated for 1 h at room temperature. After DPBS washes, cell nuclei were counterstained with Hoechst 33258 (1:10,000) and lipid droplets labeled with LipidSpot 610 (1:1,000). Cells were imaged immediately after immunofluorescence staining using a Nikon CSU-W1 Spinning Disk microscope with a Nikon SR HR Plan Apo Lambda S 100X silicone immersion objective lens (NA 1.35). For co-localization analyses in 3D, Z-stacks with 0.2 μm thick slices were acquired in 1024x1024 mode in the Nikon software NIS-Elements. The 3D Volume View tool on Nikon’s GA3 in NIS-Elements was used for 3D rendering and segmentation of lipid droplets and proteins of interest (*i.e.*, HSD17B8, DHRSX and SQSTM1). The lipid droplet volume interacting with the protein volume was extracted and divided over the total lipid droplet volume quantified in each image.

### BODIPY-C_12_ pulse-chase and starvation experiments

iPSC-derived astrocytes plated on 96-well μ-Plates (Ibidi 89626) were labeled for 2 h with 0.5 μM BODIPY FL C_12_ lipid (Thermo Fisher Scientific D3822) diluted in AM media without FBS. Cells were then washed three times and incubated 2 h in AM media without FBS to allow for the fluorescent lipids to incorporate into lipid droplets. Following this incubation, cells were chased in AM media without FBS and live imaged at 18h and 24h timepoints. Lipid droplets were labeled with LipidSpot (1:1,000) (Biotium) and cytosol was labeled with ViaFluor dye (1:5,000) (Biotium). Both dyes were present across the chase period.

To assess lipid droplet dynamics under autophagy-inducing conditions (starvation), astrocytes were starved by applying HBSS (Thermo Fisher Scientific 14025082) as cell culture medium for 1h (Figure 5F). Cells were then fixed for immunofluorescence analysis. To measure lipid droplet accumulation in the context of prolonged starvation, we cultured astrocytes in AM media containing FBS without media changes for 5 days (Figure S5D).

### Statistics and data analysis

Details of analyses can be found in figure legends. All statistical analysis for image quantification and functional assays was performed using GraphPad Prism.

## Contributions

C.C-L., R.vd.K., M.G. and P.S.N. conceived the project. C.C-L., J.T.R., Y.H., I.K., N.B., R.G., L.G.Y. and P.S.N. developed the methodology. Data analysis was performed by C.C-L., J.T.R., Y.H., I.K., N.B., S.J.K-H. and P.S.N. C.C-L. validated the findings. C.C-L., J.T.R., S.J.K-H. and P.S.N. were responsible for data visualization. C.C-L. and P.S.N. wrote the original draft, and all authors contributed to reviewing and editing the manuscript. Resources were provided by C.C-L., J.T.R., Y.H., I.K., N.B., R.G., L.G.Y., S.J.K-H., M.R.C., R.vd.K., M.G., Y.A.Q. and P.S.N. Supervision was provided by M.R.C., R.vd.K., M.G., Y.A.Q. and P.S.N. Funding was acquired by M.R.C., R.vd.K., M.G., Y.A.Q. and P.S.N.

## Acknowledgements

We would like to thank Dr. Marianita Santiana for helpful guidance on confocal imaging and 3D analysis. We thank members of the Narayan Lab, as well as Dr. Richard Proia, Dr. Ashley Frakes, Dr. Richard Youle, Dr. Michael Ward, Dr. Sonali Mohan, and Dr. Lance Johnson for helpful discussions of this work. Figure 1A was created using Biorender (https://www.biorender.com).

This research was supported by the Intramural Research Programs of the National Institute of Diabetes and Digestive and Kidney Diseases (NIDDK) (1ZIADK075158) and National Institute of Aging within the National Institutes of Health (NIH). This work was also funded by the Chan Zuckerberg Initiative, collaborative pairs grant DAF2022-250616 (R.vd.K. and M.G.) and a Vidi 2021 (09150172110086) from the Dutch Research Council (NWO) grant (R.vd.K.).

The contributions of the NIH authors (C.C-L., J.T.R., Y.H., I.K., R.G., L.G.Y., M.R.C., Y.A.Q. and P.S.N) were made as part of their official duties as NIH federal employees, are in compliance with agency policy requirements, and are considered Works of the United States Government. However, the findings and conclusions presented in this paper are those of the authors and do not necessarily reflect the views of the NIH or the U.S. Department of Health and Human Services.

## Data availability

Raw proteomics data generated during this study will be deposited on ProteomeXchange and will be available upon publication. Lipidomics data will be available via the Neurolipid Atlas website upon publication.

## Supplementary Figures

**Figure S1.**
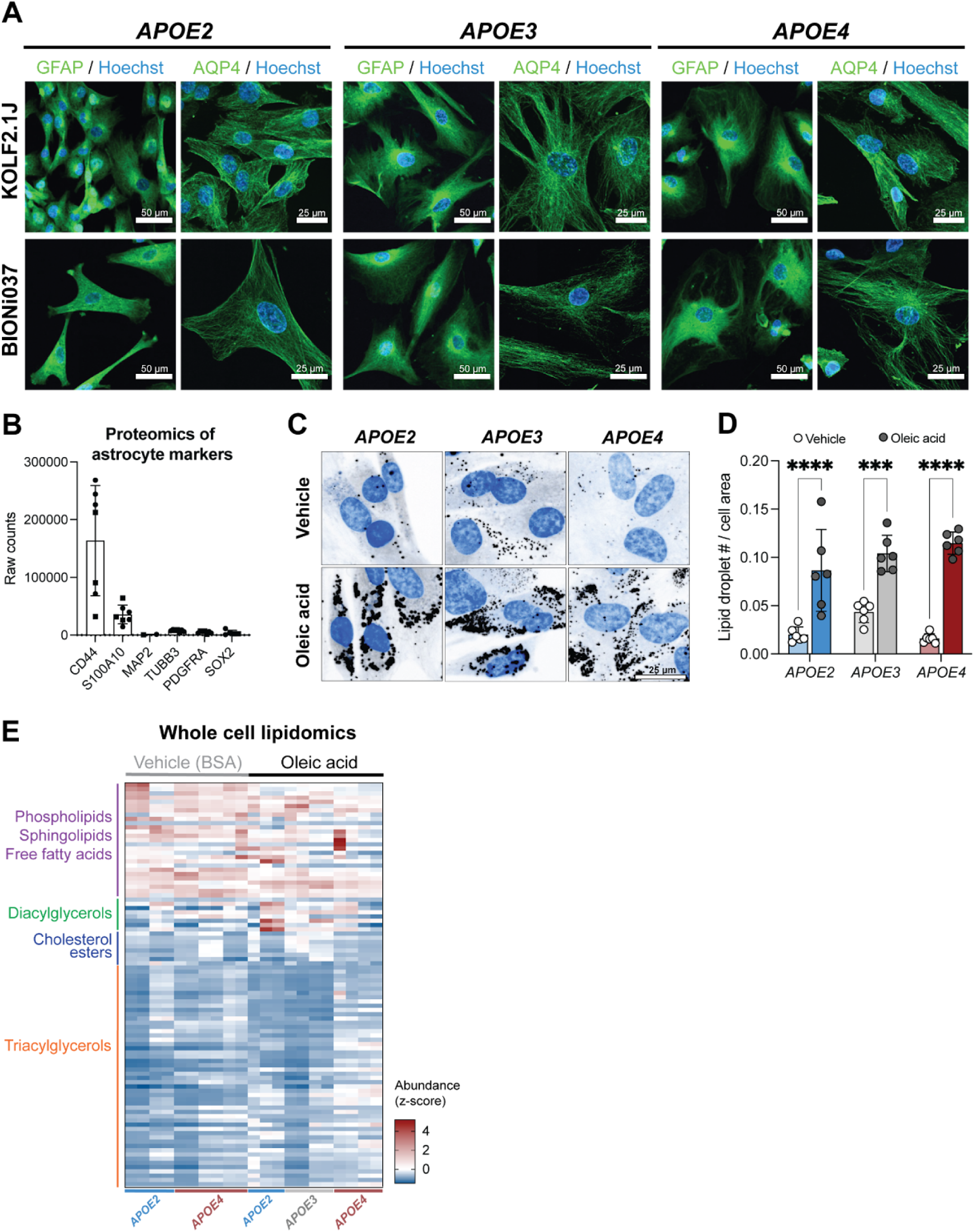
Data relating to Figure 1. A. iPSC-derived astrocytes from KOLF2.1J (top) and BIONi037 (bottom) isogenic series both stain for the canonical astrocytic markers, GFAP and AQP4. Scale bars are 25 μm for AQP4 and 50 μm for GFAP. B. Raw abundance counts of protein markers for astrocytes (CD44 and S100A10) and non-astrocytic marker proteins (neurons–MAP2, TUBB3; oligodendrocytes–PDGFRA; and iPSCs–SOX2). Measurements were derived from whole cell proteomics data. Square symbols represent KOLF2.1J-derived lines and circles represent BIONi037-derived lines. Each point represents data from an independent line and derivation, n=3-4. C. Immunofluorescence of neutral lipids (LipidSpot, black) in isogenic iPSC-derived astrocytes (BIONi037-derived) homozygous for three common *APOE* genotypes in the presence of either vehicle (BSA) or 80 μM oleic acid. Blue staining is Hoechst 33258 (nuclei). Scale bar is 25 μm. D. Quantification of lipid droplets per cell area in isogenic iPSC-derived astrocytes (BIONi037-derived) homozygous for three common *APOE* genotypes in the presence of either vehicle (BSA) or 80 μM oleic acid. Data represent n=6 independent treatments, represented as mean ± SD. *** *P* ≤ 0.001,**** *P* ≤ 0.0001 by two-way ANOVA with post-hoc Šídák’s test. E. A heatmap depicting mass spectrometry lipidomics measurements (z-score of abundance normalized to total lipid content) on unfractionated “Whole Cell” samples when cells are treated with vehicle (BSA) or oleic acid (80 μM). N=10-11 independent cell growths across two isogenic sets of human iPSC-derived astrocytes are shown with representation from multiple lipid classes.

**Figure S2.**
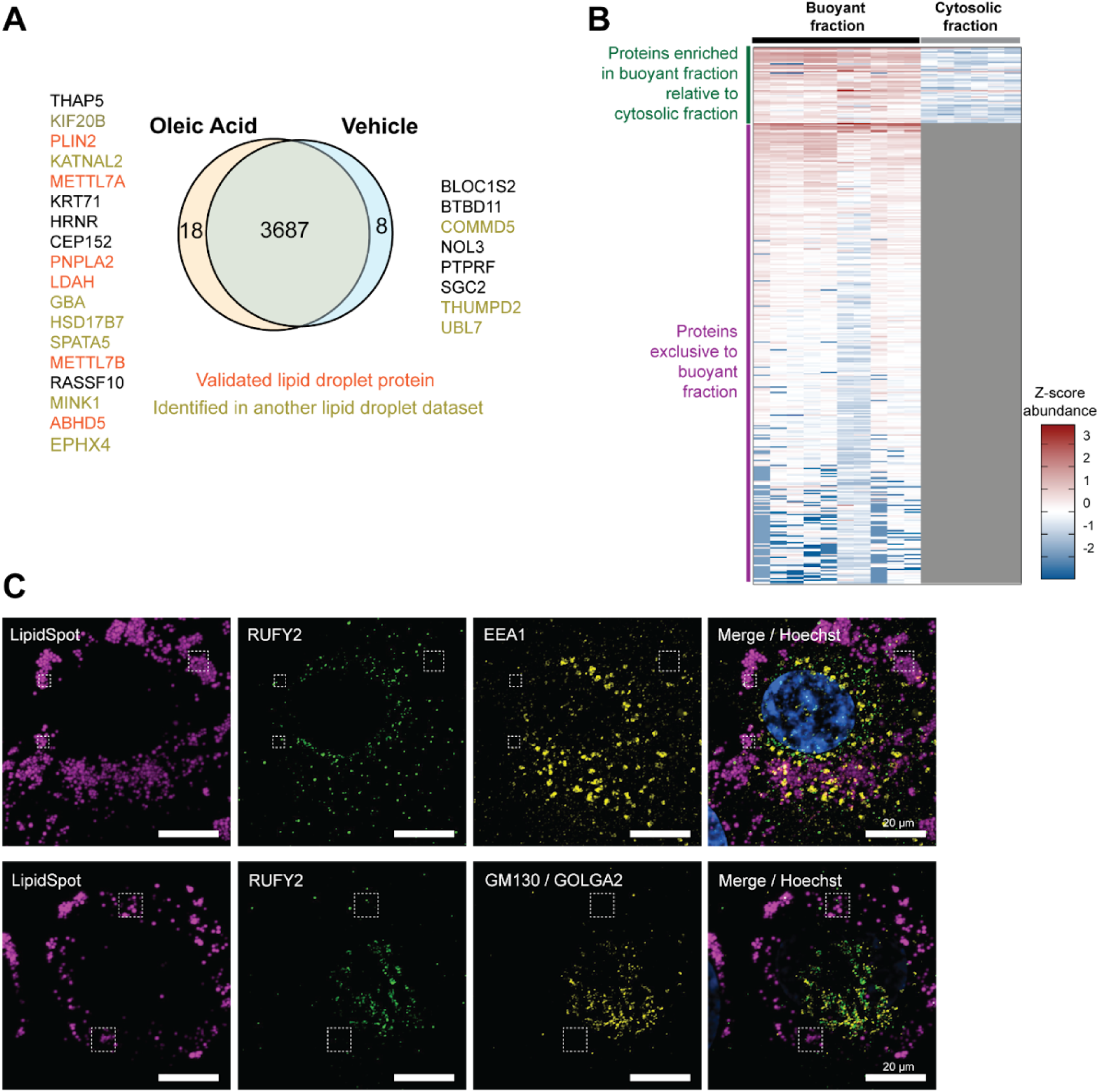
Data relating to Figure 2. A. Venn diagram comparing proteins present in the buoyant fractions of BSA-(vehicle) and oleic acid-treated human iPSC-derived astrocytes. Only 26 proteins were significantly differentially represented in these two treatment conditions. Orange proteins are canonical lipid droplet proteins validated by immunofluorescence in other studies. Yellow-green represents proteins identified in other published lipid droplet proteome studies from other cell types. B. A heatmap depicting the mass spectrometry proteomics data (z-score of abundance) of proteins selected as part of our human iPSC-derived astrocyte lipid droplet-associated proteome in the buoyant fraction samples (n=10) and the cytosolic fractions (n=6). C. Immunofluorescence staining of RUFY2 (green), lipid droplets (magenta), early endosomes (EEA1, yellow), golgi (GM130/GOLGA2; yellow) and nuclei (blue). Dashed squares show areas where RUFY2 is localized to lipid droplets but not to early endosomes or golgi. Scale bars are 20 μm.

**Figure S3.**
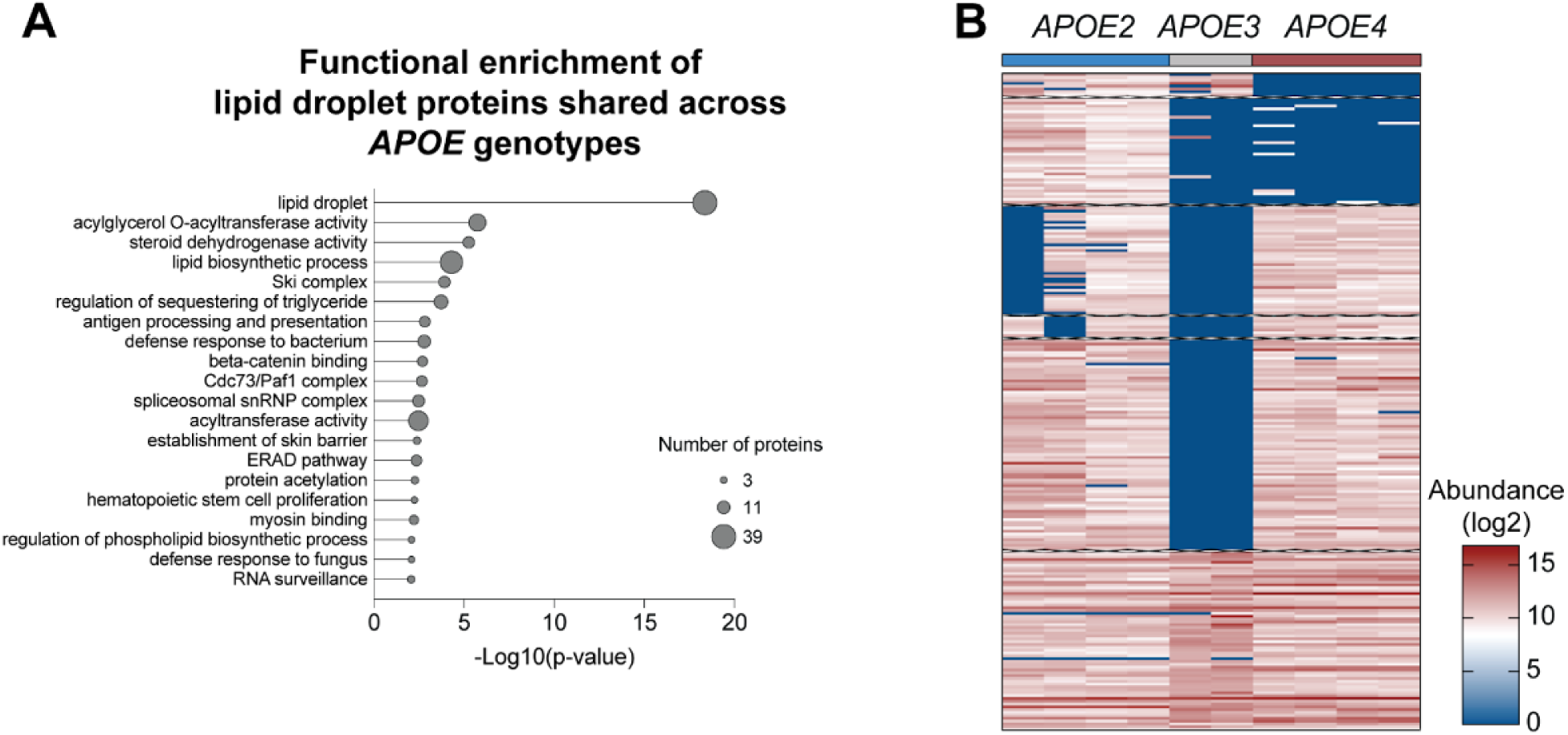
Data relating to Figure 3. A. Overrepresentation analysis to determine functional enrichment of lipid droplet-associated proteins commonly present in the lipid droplet-associated proteomes of all three *APOE* genotypes. B. A heatmap depicting the mass spectrometry proteomics data (z-score of abundance) of proteins significantly differentially abundant in lipid droplets from iPSC-derived astrocytes harboring different *APOE* genotypes.

**Figure S4.**
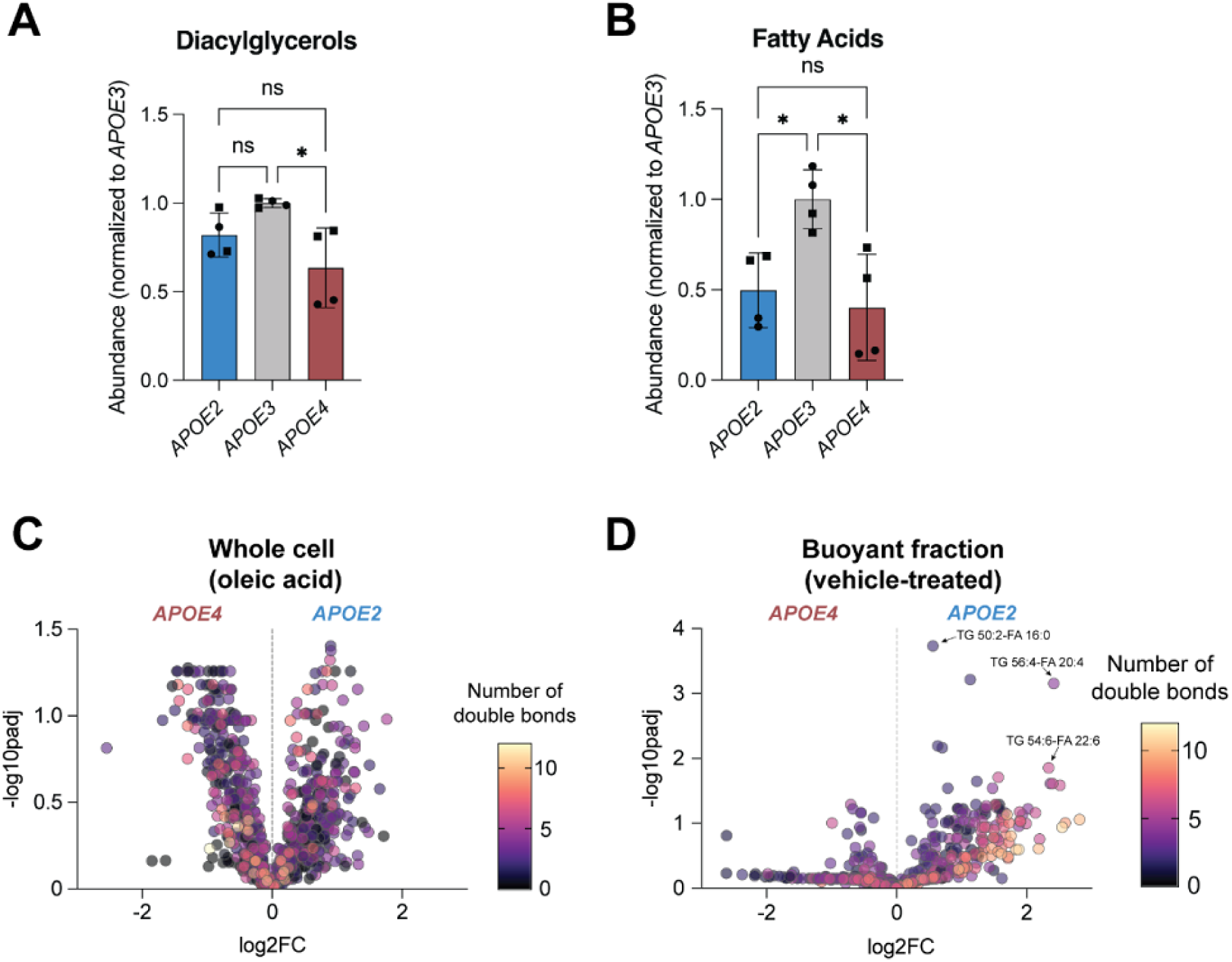
Data relating to Figure 4. A. Abundance of diacylglycerols in buoyant fractions of various *APOE* genotypes obtained from iPSC-derived astrocytes treated with oleic acid. Data represent n=4 independent replicates across two isogenic sets of human iPSC-derived astrocytes. * *P* ≤ 0.05 by one-way ANOVA with post-hoc Tukey’s test. B. Abundance of free fatty acids in buoyant fractions of various *APOE* genotypes obtained from iPSC-derived astrocytes treated with oleic acid. Data represent n=4 independent replicates across two isogenic sets of human iPSC-derived astrocytes. * *P* ≤ 0.05 by one-way ANOVA with post-hoc Tukey’s test. C. Volcano plots comparing whole cell lipids between *APOE4* and *APOE2* iPSC-derived astrocytes following treatment with oleic acid. Each data point represents individual lipid species from n=4 independent replicates across two isogenic sets of human iPSC-derived astrocytes. Color coding represents fatty acid saturation (number of double bonds). D. Volcano plots comparing buoyant fraction lipids between *APOE4* and *APOE2* iPSC-derived astrocytes following treatment with vehicle (BSA). Each data point represents individual lipid species from n=4 independent replicates across two isogenic sets of human iPSC-derived astrocytes. Color coding represents fatty acid saturation (number of double bonds).

**Figure S5.**
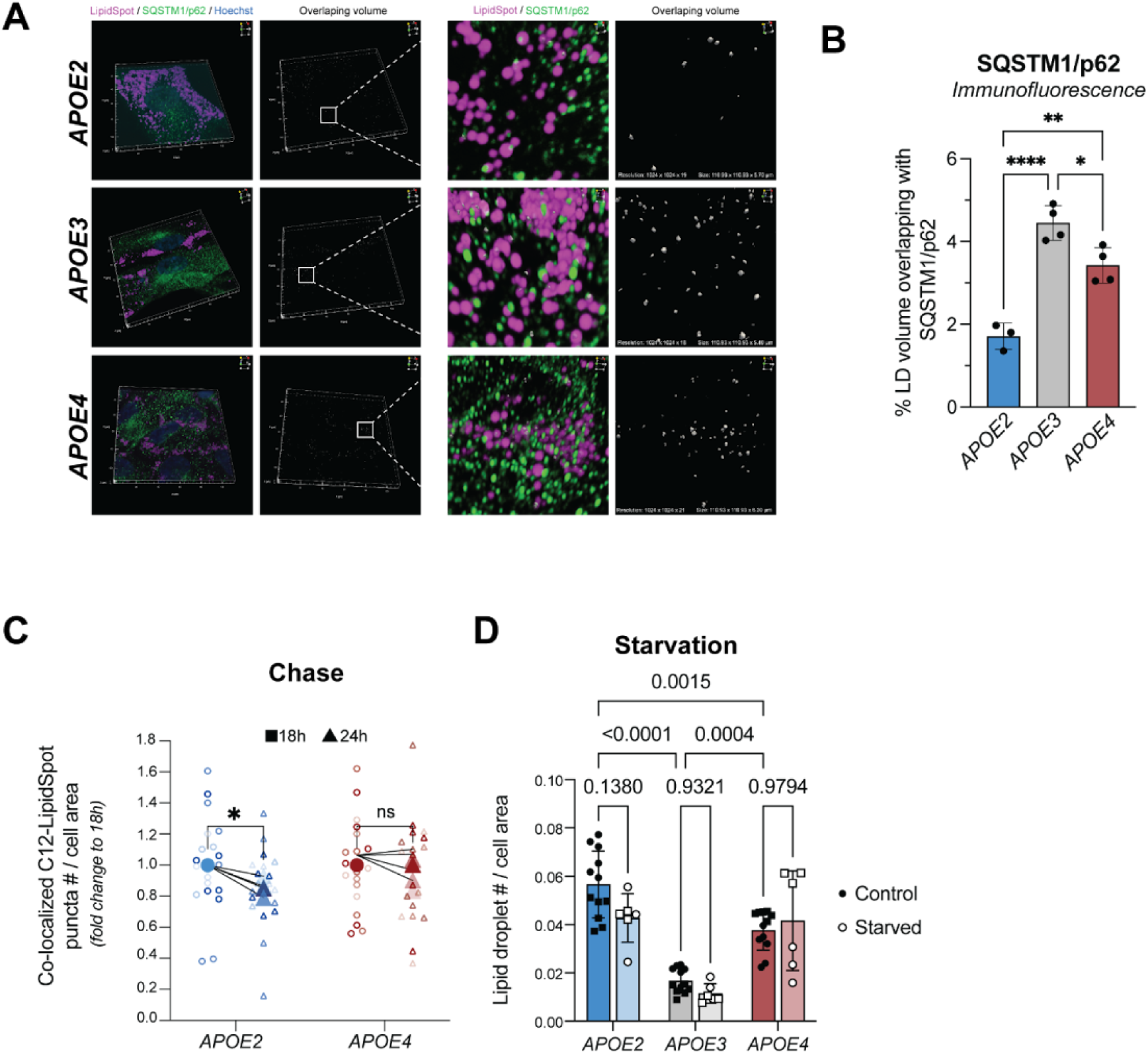
Data relating to Figure 5. A. Immunofluorescence staining of SQSTM1/p62 in green that contacts lipid droplets (magenta) in human iPSC-derived astrocytes harboring the three *APOE* genotypes (shown in BIONi037 line). Overlap volume is shown in white. Scale of 3D projection shown is 110.93 x 110.93 x 10-6.30 μm^3^. B. Quantification of percent of lipid droplet volume overlapping with SQSTM1/p62 signal in 3D confocal imaging (shown in BIONi037 line). Data represent quantification of an average of ∼700 lipid droplets per imaging frame. Data plotted are n=3-4 imaging frames per *APOE* genotype. Data are represented as mean ± SD. **** *P* ≤ 0.0001, ** *P* ≤ 0.01, * *P* ≤ 0.05 by one-way ANOVA with post-hoc Tukey’s test. C. The change in number of BODIPY-C_12_-positive lipid droplets (co-labeled with LipidSpot) during a chase period of 18 h and 24 h in human iPSC-derived astrocytes harboring *APOE2* and *APOE4* genotypes (shown in BIONi037 line). Data shown in large symbols represent n=4 independent pulse chase experiments calculated as fold change to 18 h. Data shown in small symbols represent individual replicates for each experiment. * *P* ≤ 0.05 by repeated measures two-way ANOVA with post-hoc Šidák’s test. D. Quantification of lipid droplets per cell area in two isogenic sets of human iPSC-derived astrocytes under regular culture conditions (control) and starvation for 5 days (starved). Each point represents an independent treatment. Data are represented as mean ± SD. **** *P* ≤ 0.0001 by two-way ANOVA with post-hoc Tukey’s test.

